# *Gabra2* is a genetic modifier of Dravet syndrome in mice

**DOI:** 10.1101/2020.04.19.048546

**Authors:** Nicole A. Hawkins, Toshihiro Nomura, Samantha Duarte, Robert W. Williams, Gregg E. Homanics, Megan K. Mulligan, Anis Contractor, Jennifer A. Kearney

## Abstract

Pathogenic variants in epilepsy genes result in a spectrum of clinical presentation, ranging from benign phenotypes to intractable epilepsies with significant co-morbidities and increased risk of sudden unexpected death in epilepsy (SUDEP). One source of this phenotypic heterogeneity is modifier genes that affect penetrance, dominance or expressivity of a primary pathogenic variant. Mouse models of epilepsy also display varying degrees of clinical severity on different genetic backgrounds. Mice with heterozygous deletion of *Scn1a* (*Scn1a*^*+/−*^) model Dravet syndrome, a severe epilepsy most often caused by *SCN1A* haploinsufficiency. *Scn1a*^*+/−*^ heterozygous mice recapitulate key features of Dravet syndrome, including febrile and afebrile spontaneous seizures, SUDEP, and cognitive and behavioral deficits. The *Scn1a*^*+/−*^ mouse model also exhibits strain-dependent phenotype severity. *Scn1a*^*+/−*^ mice maintained on the 129S6/SvEvTac (129) strain have normal lifespan and no overt seizures. In contrast, admixture with C57BL/6J (B6) results in severe epilepsy and premature lethality in [B6×129]F1.*Scn1a*^*+/−*^ mice. In previous work, we identified Dravet Survival Modifier loci (*Dsm1-Dsm5*) responsible for strain-dependent differences in survival. *Gabra2*, encoding the GABA_A_ α2 subunit, was nominated as the top candidate modifier at the *Dsm1* locus on chromosome 5. Direct measurement of GABA_A_ receptors found lower abundance of α2-containing receptors in hippocampal synapses of B6 mice relative to 129. We also identified a B6-specific single nucleotide intronic deletion within *Gabra2* that lowers mRNA and protein by nearly 50%. Repair of this *de novo* deletion reestablished normal levels of *Gabra2* transcript and protein expression. In the current study, we used B6 mice with the repaired *Gabra2* allele to validate it as a modifier of phenotype severity in *Scn1a*^*+/−*^ mice. Repair of *Gabra2* restored transcript and protein expression, increased abundance of α2-containing GABA_A_ receptors in hippocampal synapses, and improved seizure and survival phenotypes of *Scn1a*^*+/−*^ mice. These findings validate *Gabra2* as a genetic modifier of Dravet syndrome.

## Introduction

Dravet syndrome is a severe, infant-onset epileptic encephalopathy with at least 80% of cases resulting from *de novo* pathogenic variants in *SCN1A* (1). Individuals with Dravet syndrome display multiple seizure types that are often refractory to standard therapeutics, an elevated SUDEP risk, cognitive and behavioral deficits, and motor system dysfunctions (2, 3). Phenotype heterogeneity is common in monogenic epilepsy syndromes, with a spectrum of clinical presentations, ranging from benign seizures to intractable epilepsy and increased SUDEP risk (4–13). Modifier genes that affect penetrance and expressivity are likely contributors to this variability.

*SCN1A* haploinsufficiency is the major cause of Dravet syndrome; therefore, mice with heterozygous deletion of *Scn1a* were developed to model Dravet syndrome (14–16). *Scn1a*^*+/−*^ mice recapitulate several core features of Dravet syndrome, including febrile and afebrile seizures, elevated risk of sudden death, and neurobehavioral and cognitive deficits (14, 17–21). These phenotypes are highly penetrant on the C57BL/6J (B6) genetic background, but are absent when the mutation is on inbred 129 strain backgrounds (14–16, 22). Taking advantage of the strain difference, we previously mapped Dravet syndrome modifier (*Dsm*) loci responsible for strain-dependent differences in survival. *Gabra2*, encoding the α2 subunit of the GABA_A_ receptor, was the highest ranked candidate gene at the *Dsm1* locus (14, 23). Notable differences in *Gabra2* expression among several inbred mouse strains had been previously reported (24–26). Further work investigating a *Gabra2* eQTL identified a single nucleotide *de novo* deletion in a splice acceptor site that was only present in B6 genomes after 1976 (25, 27). This spontaneous mutation causes a global reduction of *Gabra2* transcript and protein expression in brain compared to older B6 strains and to 16 other common and wild-derived inbred mouse strains (27). Repair of the single nucleotide deletion within the *Gabra2* intron fully restores expression back to the level of other inbred strains (27). However, it is not clear whether this nucleotide variant contributes to the strain-dependent phenotype differences observed in *Scn1a*^*+/−*^ Dravet mice.

In the current study, we sought to determine if this single nucleotide deletion was responsible for the *Dsm1* modifier effect. We used B6 mice carrying a repaired (edited) *Gabra2* allele (Edited/B6 mice) and crossed them with 129.*Scn1a*^*+/−*^ mice to ascertain effects on Dravet phenotypes. We observed rescue of seizure, survival and neuronal phenotypes with this single nucleotide repair, confirming *Gabra2* as an epilepsy modifier gene.

## Results

In prior genetic mapping studies, we identified *Gabra2* as the top *Dsm1* candidate modifier gene influencing survival of *Scn1a*^*+/−*^ mice (23). We proposed that differential expression of *Gabra2* between 129 and B6 as the likely mechanism. In parallel work, we attributed lower expression in B6 to a single nucleotide intronic deletion present only in the current B6 genome. Repair of this deletion by re-insertion of the wild-type nucleotide via CRISPR/Cas9 editing restored expression of *Gabra2* to again match that of other mouse strains (Figure 1A) (27). The B6 mice with restored expression of *Gabra2* (Edited) provide a definitive resource to test the hypothesis that strain-dependent differences in *Gabra2* expression are responsible for the *Dsm1* modifier effect and have even enabled us to localize the effect to specific nucleotide differences among the parental strains.

**Figure 1.**
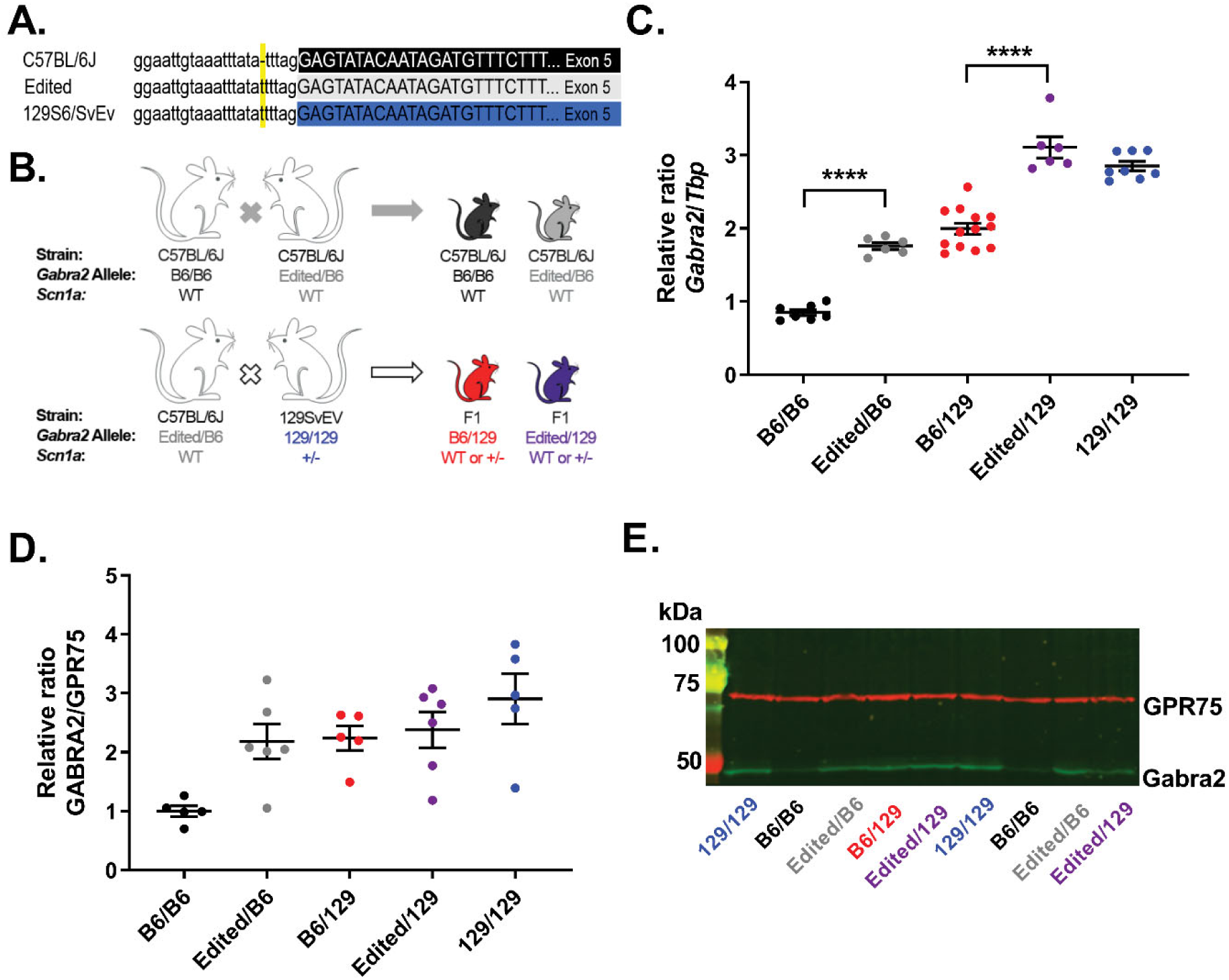
Editing of *Gabra2* B6 allele normalizes expression relative to 129. **A)** The B6 *Gabra2* intronic deletion is located on Chr 5 at 71,014,638 bp (GRCm38.p6) (yellow). Sequence of the B6 allele (black) is compared to the Edited (grey) and 129S6/SvEvTac alleles (blue). **B)** Breeding scheme for mice used in the study. For expression and electrophysiology experiments, isogenic B6.*Gabra2* Edited mice were crossed with B6 to generate offspring with Edited/B6 (grey) or B6/B6 (black) alleles at *Gabra2*. For expression and seizure experiments B6.*Gabra2* mice were crossed with isogenic 129.*Scn1a*^*+/−*^ mice to generate F1*.Scn1a^+/−^* or F1 *Scn1a*^+/+^ mice with Edited/129 (purple) or B6/129 (red) alleles at *Gabra2*. **C)** Relative expression of *Gabra2* transcript assayed by quantitative RT-ddPCR on whole brain samples from mice with B6/B6, Edited/B6, B6/129, Edited/129 and 129/129 alleles at *Gabra2*. Transcript expression differed between genotypes (F_4,35_=104.5, p<0.0001; one-way ANOVA). Allele-specific expression differences were detected for samples with B6/B6 (0.85±0.04), B6/129 (2.0±0.1) and 129/129 (2.9±0.1) alleles at *Gabra2* (p<0.0001, Tukey’s; significance not indicated on graph). Expression differed between Edited/B6 (1.8±0.1) and Edited/129 (3.1±0.1) alleles compared to unedited littermate controls. **** p<0.0001, Tukey’s. Symbols represent samples from individual mice, horizontal lines represent group averages, and error bars represent SEM with 6-13 mice per genotype. **D)** Quantification of protein expression determined from western blots. Relative expression of whole brain GABRA2 protein in samples with B6/B6 (1.0±0.1), Edited/B6 (2.2±0.3), B6/129 (2.2±0.2), Edited/129 (2.4±0.3) and 129/129 (2.9±0.4) alleles at *Gabra2* assayed by western blot. Relative GABRA2 protein differed between genotypes (F_4,22_=5.349, p=0.0037; One-way ANOVA). Allele-specific expression differences were detected for samples with B6/B6 and 129/129 alleles at *Gabra2* (p<0.0017, Tukey’s; significance not indicated on graph). Symbols represent samples from individual mice, horizontal lines represent group averages, and error bars represent SEM with 5-6 mice per genotype. **E)** Representative western blot.

### Single nucleotide repair of *Gabra2* increases transcript and protein expression

We first confirmed that repair of *Gabra2* (Edited/B6) altered allele-specific expression when crossed with the 129 strain (Figure 1B). We evaluated transcript and protein expression in wild-type (WT) mice carrying the following alleles at *Gabra2*: B6/B6, Edited/B6, B6/129, Edited/129 or 129/129 (Figure 1B). *Gabra2* transcript expression differed between genotypes (F_4,35_=104.5, p<0.0001, One-way ANOVA) (Figure 1C). Allele-specific expression differences between B6/B6, 129/B6 and 129/129 were similar to prior reports (23). Expression of *Gabra2* transcript in Edited/B6 did not differ from B6/129, while both were elevated relative to B6/B6 (Figure 1C) (p<0.0001, Tukey’s). *Gabra2* expression levels in Edited/129 were similar to 129/129, while expression in B6/129 was lower (Figure 1C) (p<0.0001, Tukey’s). GABRA2 protein expression followed the same pattern as transcript and differed between genotypes, (F_4,22_=5.349, p=0.0037, One-way ANOVA) (Figure 1D/E). Average GABRA2 expression in B6/B6 was approximately 3-fold lower relative to 129/129 mice (p<0.0017, Tukey’s). GABRA2 expression in Edited/B6 was similar to B6/129 (Figure 1D). These results demonstrate that repair of the B6 *Gabra2* allele normalized transcript and protein expression to levels that were comparable to 129, as expected.

### Single nucleotide repair of *Gabra2* alters neuronal phenotype

We previously demonstrated that perisomatic inhibitory synapses of hippocampal CA1 neurons had a greater abundance of α2-containing GABA_A_ receptors in 129 mice compared to B6 mice [28(28). α2 GABA_A_ receptor mediated currents can be distinguished by the use of the selective α_2_/α_3_ positive allosteric modulator (PAM) AZD7325, which has a larger effect on slowing the decay kinetics of inhibitory postsynaptic currents (IPSCs) enriched in α2 subunits (28). To determine whether the Edited allele affected synaptic GABA_A_ receptors, we recorded perisomatic IPSCs in CA1 neurons in B6/B6 and Edited/B6 mice. Evoked IPSCs were desynchronized by application of extracellular Sr^2+^ so that asynchronous quantal GABAergic events (aIPSCs) could be recorded. aIPSCs from perisomatic synapses were recorded by stimulating in Stratum Pyramidale (SP) during a control period and after application of AZD7325 (100nM) and current decay times measured (Figure 2). aIPSCs from B6/B6 mice exhibited an average baseline decay of 7.07±0.28 ms during the control period, and aIPSCs from Edited/B6 mice had an average decay of 6.89±0.51 ms during the baseline period. AZD7325 application prolonged aIPSC decay times in slices from both B6/B6 mice (8.76±0.35 ms, p=0.0002, Wilcoxon) and Edited/B6 mice (9.576±0.64 ms, p=0.0010, Wilcoxon). However, the effect of AZD7325 was significantly greater in recordings from Edited/B6 slices, which exhibited a 140±5.3% increase compared to B6/B6 which had a125±2.6% increase (Figure 2D, p=0.047, Mann-Whitney). This suggests that perisomatic CA1 GABAergic synapses in Edited/B6 mice are enriched in α2-containing receptors compared to those in B6/B6 mice. As demonstrated previously AZD7325 did not affect aIPSC amplitude in either strain (B6/B6: 102 ± 3.6% and Edited/B6: 100.6 ± 4.2%, p=0.73, Mann-Whitney) (Figure 2C).

**Figure 2.**
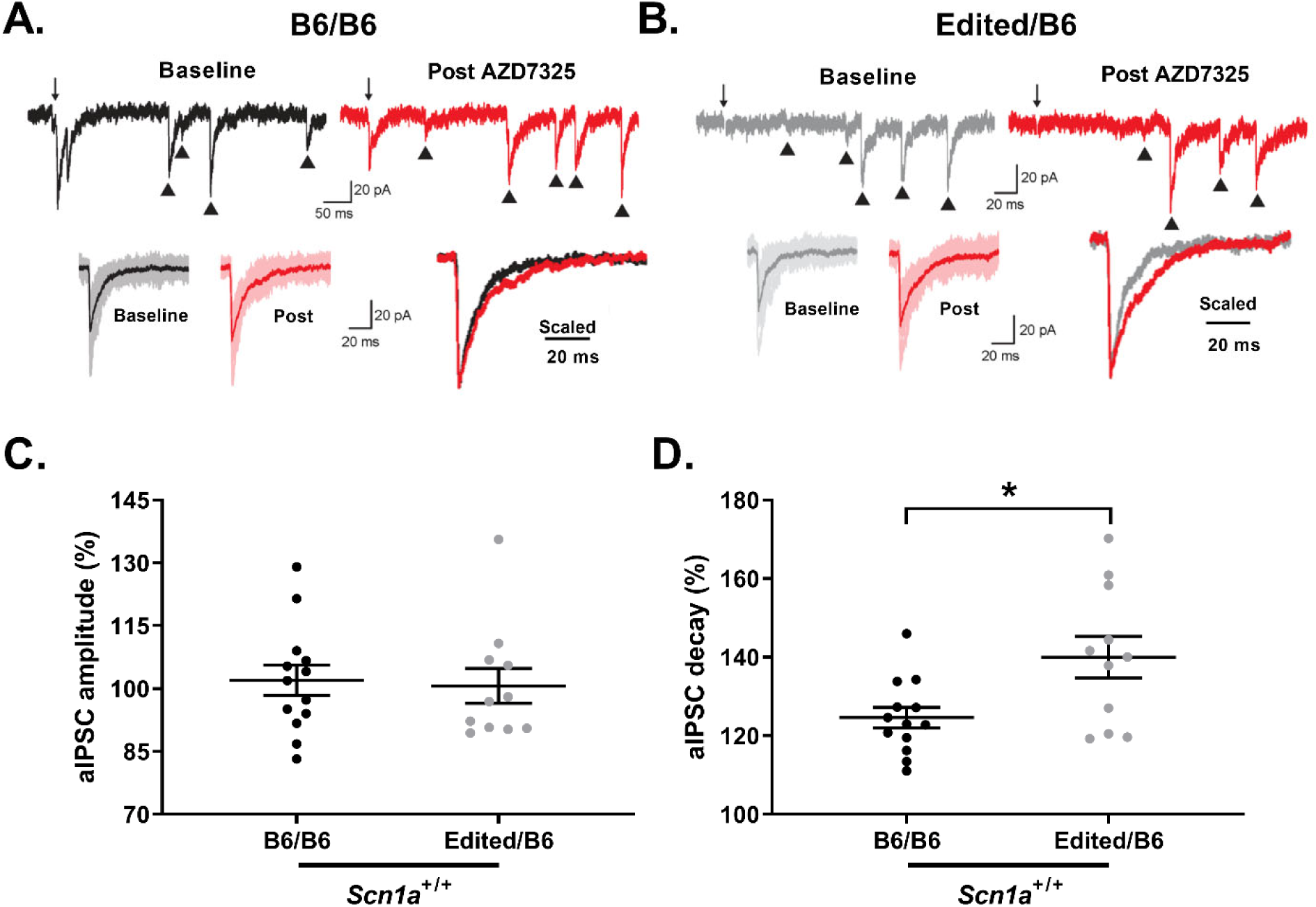
AZD7325 has a larger effect on inhibitory synapses in Edited/B6 mice. **A)** Representative traces of perisomatic aIPSCs in CA1 neurons before (baseline) and after AZD7325 (post) treatment in slices from B6/B6 mice and **B)** slices from Edited/B6 mice. **C)** Effect of AZD7325 on amplitudes of perisomatic aIPSC in CA1 of B6/B6 and Edited/B6 mice. **D)** Effect of AZD7325 on the decay kinetics of aIPSC in B6/B6 and Edited/B6 mice. AZD7325 had a greater effect on decay kinetics in Edited/B6 mice (140 ± 5.3%) compared to B6/B6 mice (125 ± 2.6%). *p=0.047, Mann-Whitney.

### Single nucleotide repair of *Gabra2* improves phenotype of F1.*Scn1a*^+/−^ mice

129.*Scn1a*^*+/−*^ mice have no overt seizure or premature lethality phenotype, whereas F1.*Scn1a*^*+/−*^ mice have spontaneous seizures and premature lethality (14–16, 22). Similarly, mapping with interval-specific congenic (ISC) strains demonstrated that homozygosity for 129 alleles in the *Gabra2* region could rescue seizure and survival phenotypes in otherwise F1.*Scn1a*^*+/−*^ mice (23). To further refine the genetic mechanism, we investigated whether repair of the B6-specific *Gabra2* intronic variant could improve survival of F1.*Scn1a*^*+/−*^ mice. We crossed B6 mice with heterozygous repair of *Gabra2* (B6/Edited) to 129.*Scn1a*^*+/−*^ to generate F1.*Scn1a*^*+/−*^ mice carrying Edited/129 or B6/129 alleles at *Gabra2* (Figure 1B). Survival was improved in F1.*Scn1a*^*+/−*^ mice with Edited/129 versus B6/129 alleles at *Gabra2* (p<0.0001, Logrank Mantel-Cox). Mice with Edited/129 alleles had 97% survival to 8 weeks of age compared to 43% of mice with B6/129 alleles (Figure 3A), supporting *Gabra2* as the modifier gene of the *Dsm1* locus.

**Figure 3.**
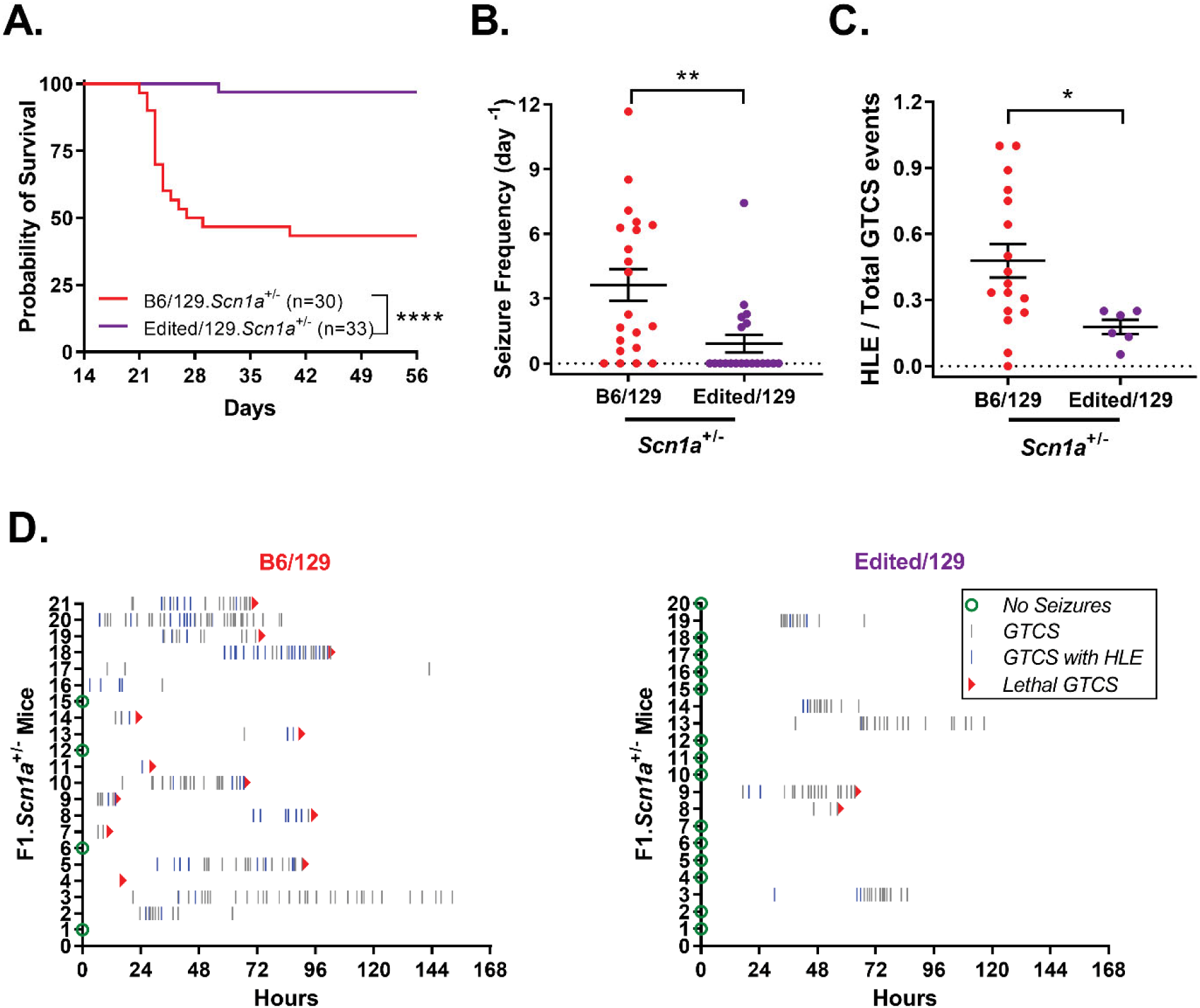
Survival and seizure burden improved in F1.*Scn1a*^*+/−*^ with Edited/129 versus B6/129 alleles at *Gabra2*. **A)** Kaplan Meier survival plot comparing 8 week survival of B6/129 and Edited/129 F1.*Scn1a*^*+/−*^ mice. Survival was worse in B6/129 mice (43%) compared to Edited/129 (97%) with n=30-33 per genotype. ****p<0.0001, Logrank Mantel-Cox. **B)** The proportion of mice exhibiting spontaneous GTCS and average GTCS frequency differed between F1.*Scn1a*^*+/−*^ mice with B6/129 (81% with seizures; 3.6±0.7 GTCS/day) versus Edited/129 (30% with seizures; 0.9±0.41 GTCS/day) alleles at *Gabra2*. Symbols represent samples from individual mice, horizontal lines represent the group average, and error bars represent SEM with n=20-21 per genotype. **p<0.0018, Mann-Whitney. **C)** Among mice with GTCS, the average proportion of GTCS that progressed to HLE differed between mice with B6/129 (0.48±0.08) versus Edited/129 (0.18±0.03) alleles at *Gabra2*. Symbols represent samples from individual mice, and error bars represent SEM with n=6-17 per genotype. **p<0.015, Mann-Whitney. **D)** Spontaneous GTCS diary plots for individual F1.*Scn1a*^*+/−*^ mice with B6/129 or Edited/129 alleles at *Gabra2*. Each row represents a single F1.*Scn1a*^*+/−*^ mouse (n=20-21 per genotype) over the 168 hour monitoring period or until occurrence of sudden, unexpected death indicated by a red triangle. Green circles represent subjects with no GTCS events. Grey tick marks indicate a GTCS without HLE. Blue tick marks represent a GTCS with HLE.

Next, we investigated if the *Gabra2* variant altered seizure frequency and/or severity. At P18-19, F1.*Scn1a*^*+/−*^ mice with Edited/129 and B6/129 alleles at *Gabra2* were subjected to a single hyperthermia-induced priming seizure and quickly cooled to baseline temperature(18). The temperature for seizure onset did not differ between Edited/129 (41.1±0.2°C) and B6/129 (41.2±0.2°C) groups (p= 0.9127, Student’s T-test), and all mice in both groups had a seizure. Following the priming seizure, mice were continuously monitored for spontaneous generalized tonic-clonic seizures (GTCS) for 7 days or until sudden death occurred. The proportion of F1.*Scn1a*^*+/−*^ exhibiting GTCS during the monitoring period differed between *Gabra2* genotypes (p<0.0026, Fisher’s exact). Only 30% of mice with Edited/129 alleles exhibited seizures, while 81% of mice with B6/129 alleles had seizures (Figure 3B/D). Seizure frequency in F1.*Scn1a*^*+/−*^ mice with Edited/129 alleles was lower (0.9±0.4 GTCS/day) relative to mice with B6/129 alleles (3.6±0.7 GTCS/day) (Figure 3B/D) (p<0.0018, Mann-Whitney). While previous studies demonstrated no correlation between seizure frequency and survival, it was reported that seizure severity, more specifically the occurrence of tonic hindlimb extension (HLE) during a GTCS, was correlated with survival (18, 29, 30). Therefore, we also assessed seizure severity based on presence or absence of HLE during each GTCS event occurring in the 7-day monitoring period. GTCS events in mice with Edited/129 alleles progressed to HLE less often (18±3% of events) relative to mice with B6/129 alleles (48±8% of events) (p<0.015, Mann-Whitney) (Figure 3C/D). Together, these data demonstrate that repair of the B6 *Gabra2* allele lessened seizure burden of F1.*Scn1a*^*+/−*^ mice, supporting *Gabra2* as the genetic modifier at *Dsm1*.

## Discussion

In the present study, we demonstrated that *Gabra2* is the genetic modifier at *Dsm1* responsible for the strain-dependent difference in survival between F1 and 129 *Scn1a*^*+/−*^ mice (14, 23). Furthermore, we defined the responsible nucleotide difference underlying the modifier effect. Editing of the B6-specific single nucleotide intronic deletion in *Gabra2* normalized brain transcript and protein expression relative to the 129 allele, elevated enrichment of α2-containing GABA_A_ receptors in hippocampal synapses, and dramatically improved seizure and survival phenotypes in the F1.*Scn1a*^*+/−*^ Dravet mouse model. This work has clear therapeutic implications and suggests that interventions that increase CNS expression or function of GABRA2 should improve outcomes in Dravet syndrome.

In previous work we and others demonstrated allele-specific expression of *Gabra2*. We also established that unusually low GABRA2 protein expression is caused by a non-coding single nucleotide deletion in the C57BL/6J genome that alters splicing efficiency (24–27, 29). B6 mice are the most commonly used laboratory mouse strain and was the first to be sequenced (GRCm38). With drastic improvements in next-generation sequencing techniques, it has become evident the GRCm38 reference genome, generated from filial (F) generation 204-207 mice, may differ from the current (~F226) B6 mouse strain (31, 32). Recently, Sarani et al sequenced C57BL/6JEve, the “mother” (F223) of the current laboratory B6 animals sourced from the Jackson Laboratory (F226) (32). The group identified 59 indels and inversions between B6-Eve and GRCm38, many located within noncoding intronic regions (32). Furthermore, recent efforts from the Jackson Laboratory identified 1083 variants which passed quality controls between the GRCm38 reference sequence and sequencing from the most recent inbreeding generations of C57BL/6J (31). These variants can provide insight into the private *de novo* mutations in the B6 genome compared to other inbred strains. This is of particular interest when investigating genetic modifiers of diseases, including epilepsy. Three additional modifier loci of the *Scn1a*^*+/−*^ phenotype (*Dsm2, Dsm3, Dsm5*) have yet to be investigated (14). Preliminary examination of these private variants may suggest additional candidate modifier genes of survival in the mouse model of Dravet syndrome.

This study also highlights the importance of continued efforts in identifying modifier genes in mouse models. Several epilepsy modifier genes, including *CACNA1G* and *GABRA2*, were first identified in the *Scn1a*^*+/−*^mouse model of Dravet syndrome and later confirmed as epilepsy risk genes in humans (14, 23, 33–44). Identifying modifier genes can provide refined insights into the molecular basis of genetic disease. Furthermore, it may provide the basis for improving predictions about disease course and clinical management. Finally, modifier genes and pathways can provide novel targets for therapeutics. Previously, we used AZD7325, a GABA_A_ α2/α3-selective PAM, to modulate the neuronal phenotype of *Scn1a*^*+/−*^ mice and showed it had protective effects against hyperthermia-induced seizures (28). The current study confirmed that AZD7325 can be used to distinguish the GABA_A_ receptor type in perisomatic synapses and directly demonstrated that the Edited allele contributes to an enriched α2 subunit content. Future studies assessing efficacy of AZD7325 treatment on spontaneous seizures and survival would provide further support for targeting α_2_-containing GABA_A_ receptors for the treatment of Dravet syndrome and, potentially other epileptic encephalopathies that share reduced GABAergic signaling as a common pathogenic mechanism.

## Materials and Methods

### Ethics Statement

All animal care and experimental procedures were approved by the Northwestern University Animal Care and Use Committee (#IS00000539) in accordance with the National Institutes of Health Guide for the Care and Use of Laboratory Animals.

### Mice

CRISPR/Cas9 editing was used to insert a single intronic nucleotide into *Gabra2* on the C57BL/6J (B6) strain, B6.*Gabra2*^*em1Geh*^, as previously described (27). Heterozygous edited *Gabra2* mice (Edited/B6) were maintained by continual backcrossing to C57BL/6J (Jackson Labs, #000664, Bar Harbor, ME).

S*cn1a*^*tm1Kea*^ mice with deletion of the first coding exon, were generated by homologous recombination in TL1 ES cells as previously described in (14). This line has been maintained by continual backcrossing of heterozygotes (abbreviated as *Scn1a*^*+/−*^) to 129S6/SvEvTac inbred mice (129, Taconic, #129SVE, Rensselaer, NY).

To generate double mutant mice, heterozygous edited *Gabra2* mice were crossed to 129.*Scn1a*^*+/−*^ mice. The resulting offspring had an overall F1 genome background with B6/129 or Edited/129 alleles at *Gabra2* and WT (+/+) or heterozygous deletion (+/−) at *Scn1a*.

Mice were maintained in a Specific Pathogen Free (SPF) barrier facility with a 14-hour light/10-hour dark cycle and access to food and water *ad libitum*. Female and male mice were used for all experiments. The principles outlined in the ARRIVE (Animal Research: Reporting of *in vivo* Experiments) guideline and Basel declaration (including the 3 R concept) were considered when planning experiments.

### Genotyping

DNA was isolated from P14 tail biopsies using the Gentra Puregene Mouse Tail Kit according to the manufacturer’s instructions (Qiagen, Valencia, CA). For *Gabra2*, approximately 250 ng of DNA was digested with BAMH1-HF (R3136, New England Biolabs, Ipswich, MA) at 37°C for 1 hour. Digested DNA was then diluted 1:1 with nuclease-free water and used for template for digital droplet PCR (ddPCR) using ddPCR Supermix for Probes (No dUTP) (Bio-Rad, Hercules, CA, USA) and a custom TaqMan SNP Genotyping Assay (Life Technologies, Carlsbad, CA) to detect the insertion (sequence available upon request). Reactions were partitioned into droplets in a QX200 droplet generator (Bio-Rad). PCR conditions were 95°C for 10 minutes, then 44 cycles of 95°C for 30 seconds and 60°C for 1 minute (ramp rate of 2°C/sec) and a final inactivation step of 98°C for 5 minutes. Following amplification, droplets were analyzed with a QX200 droplet reader with QuantaSoft v1.6.6 software (Bio-Rad). For *Scn1a*, the genotype was determined by multiplex PCR as previously described (14).

### Transcript analysis

Whole brain RNA was extracted from mice with the following *Gabra2* alleles: B6/B6, Edited/B6, B6/129, Edited/129, and 129/129. Total RNA was isolated using TRIzol reagent according to the manufacturer’s instructions. First-strand cDNA was synthesized from 2 micrograms of RNA using oligo(dT) primer and Superscript IV reverse transcriptase (RT) according to the manufacturer’s instructions (Life Technologies). First-strand cDNA samples were diluted 1:10 and 5 μl was used as template. Quantitative ddPCR was performed using ddPCR Supermix for Probes (No dUTP) (Bio-Rad) and TaqMan Gene Expression Assays (Life Technologies) for mouse *Gabra2* (FAM-MGB-Mm00433435_m1) and *Tbp* (VIC-MGB-Mm00446971_m1) as a normalization standard. Reactions were partitioned into a QX200 droplet generator (Bio-Rad). Thermocycling conditions and analysis were performed as described for genotyping. Both assays lacked detectable signal in no-RT and no template controls. Relative transcript levels were expressed as a ratio of *Gabra2* to *Tbp* concentrations, with 6-13 biological replicates per group. Mice ranged in age from P23-41. Statistical comparison between groups was made using ANOVA with Tukey’s post-hoc comparisons (GraphPad Prism, San Diego, CA). Data are presented as mean ± SEM.

### Western blot analysis

Whole brain membrane proteins were isolated from mice with the following *Gabra2* alleles: B6/B6, Edited/B6, B6/129, Edited/129 and 129/129 mice. Membrane fractions 50 μg/lane were separated on a 7.5% SDS-PAGE gel and transferred to nitrocellulose. Blots were probed with rabbit polyclonal *Gabra2* antibody (822-GA2CL, PhosphoSolutions; RRID:AB_2492101; 1:1000) and mouse monoclonal anti-mortalin/GRP75 antibody (NeuroMab N52A/42; RRID:2120479; 1 μg/mL) which served as a normalization control. Anti-rabbit Alexa Fluor 790 and anti-mouse 680 antibodies (Jackson ImmunoResearch, 1:20,000) were used to detect signal on an Odyssey imaging system (Li-COR). Relative protein levels were determined by densitometry with Image Studio (Li-COR) and expressed as a ratio of *Gabra2* to GRP75, with 5-6 biological replicates per group. Mice ranged in age from P21-42. Statistical comparison between groups was made using one-way ANOVA with Tukey’s post-hoc comparisons (GraphPad Prism). Data are presented as mean ± SEM.

### Seizure phenotyping

*Scn1a*^*+/−*^ littermate mice carrying B6/129 or Edited/129 *Gabra2* alleles were monitored for spontaneous generalized tonic-clonic seizures (GTCS), as previously described (18). Briefly, at P18 or P19, mice were subjected to a single, hyperthermia induced GTCS and then immediately cooled to baseline temperature. If a GTCS did not occur, the mouse was excluded from the study (<1%). Two to three mice of mixed genotype and sex were placed in a monitoring cage with *ad libitium* access to standard rodent chow and water. Spontaneous GTCS frequency was captured by continuous video monitoring as previously described (18, 23). Mice were monitored beginning at midnight (12-16 hours post priming) for 7 consecutive days (168 hours). Videos were scored offline by reviewers blinded to genotype to determine the frequency and severity of spontaneous GTCS. GTCS were defined by the following minimal criteria: bilateral forelimb clonus with rearing and paddling; and the majority of events included loss of posture, as well as wild running/jumping with or without tonic hindlimb extension (HLE; hindlimbs extend at 180° relative to torso). The total number of seizures for each mouse was divided by the total hours monitored and then converted to a seizure frequency per 24 hours. The proportion of seizures with HLE was determined for each mouse based on presence or absence of tonic HLE phase for each GTCS event. Seizure frequency and severity were first compared between sexes within genotypes. No sex difference was detected; therefore, groups were collapsed across sex for analysis of genotype effect. Groups were compared between genotypes using Mann Whitney U-Tests (GraphPad Prism). Data are presented as means ± SEM.

### 8-week survival monitoring

*Scn1a*^*+/−*^ littermates with B6/129 or Edited/129 alleles at *Gabra2* were weaned into standard vivarium holding cages containing 4–5 mice of the same age and sex. Survival was monitored until 8 weeks of age. During that time, all mice were monitored daily for general health. Any mouse that was visibly unhealthy (e.g. underweight, poorly groomed, dehydrated, or immobile) was excluded from the study. All recorded deaths were sudden and unexpected, occurring in otherwise healthy appearing animals. Survival was first compared between sexes within genotypes. No sex difference was detected; therefore, groups were collapsed across sex for analysis of genotype effect. Survival statistics were calculated using time-to-event analysis with LogRank Mantel-Cox test (GraphPad Prism).

### Electrophysiology

Acute horizontal hippocampal slices (350 µm) were prepared from 22-31-day old male and female littermate mice with B6/B6 (n=3 mice) or B6/Edited (n=3 mice) alleles at *Gabra2*. Mice for this experiment were all wild-type at *Scn1a*. Brains were quickly removed after decapitation and sections were made in ice-cold sucrose-slicing artificial cerebrospinal fluid (ACSF) containing the following (in mM): 85 NaCl, 2.5 KCl, 1.25 NaH_2_PO_4_, 25 NaHCO_3_, 25 glucose, 75 sucrose, 0.5 CaCl_2_ and 4 MgCl_2_ with 10 µM DL-APV and 100 µM kynurenate on a Leica Vibratome. Slices were incubated in the same sucrose ACSF for ∼30 min at 30 °C. The solution was gradually exchanged for a recovery ACSF containing the following (in mM): 125 NaCl, 2.4 KCl, 1.2 NaH_2_PO_4_, 25 NaHCO_3_, 25 glucose, 1 CaCl_2_, and 2 MgCl_2_ with 10 µM DL-APV and 100 µM kynurenate at room temperature. Slices were transferred to a recording chamber after a recovery period of at least 1.5 hours. Recordings were made from CA1 pyramidal neurons in the hippocampus. Recording electrodes had tip resistances of 3 – 5 MΩ when filled with internal recording solution containing the following (in mM): 135 CsCl, 20 HEPES, 2 EGTA, 2 Mg-ATP, 0.5 Na-GTP, and 10 QX-314 (pH 7.25). Asynchronous IPSCs (aIPSCs) were recorded from perisomatic synapses where α_2_ GABA_A_ receptor subunits are enriched (28, 45). Perisomatic aIPSCs were recorded in voltage clamp mode (−70 mV) in the recording strontium based ACSF containing the following (in mM): 125 NaCl, 2.4 KCl, 1.2 NaH_2_PO_4_, 25 NaHCO_3_, 25 glucose, 6 SrCl_2_, 0.5 CaCl_2_, and 2 MgCl_2_ with blockers of excitatory responses, CNQX (10 µM) and D-APV (50 µM), equilibrated with 95% O_2_ and 5% CO_2_. Perisomatic aIPSCs were evoked by electrical stimuli given through monopolar extracellular stimulating electrodes filled with ACSF and placed in stratum pyramidale and were analyzed within a 50-500 ms window following the stimulus artifact (28, 45–47). Access resistance (Ra) was continuously monitored and experiments were omitted if the R_a_ changed > 20 % during the recordings. 4-5 cells were recorded per B6/B6 mouse. 3-5 cells were recorded per Edited/B6 mouse. Statistical comparison between genotype was made using Mann Whitney U-Test (GraphPad Prism). Data are presented as mean ± SEM.

## Acknowledgements

We thank Tyler Thaxton and Alexandra Huffman for technical support.

## Author Contributions

Conceptualization: MM, RW, GH, NAH, AC, JAK

Formal analysis: NAH, JAK

Funding acquisition: JAK

Investigation: NAH, SD, TN

Methodology: NAH, TN, AC, JAK

Project administration: NAH, JAK

Resources: GH

Supervision: JAK

Visualization: NAH, TN, AC, JAK

Writing – original draft: NAH, TN, JAK

Writing – review & editing: NAH, TN, SD, MM, RW, GH, AC, JAK.

